# Meta-analysis of the human upper respiratory tract microbiome reveals robust taxonomic associations with health and disease

**DOI:** 10.1101/2023.08.10.552808

**Authors:** Nick Quinn-Bohmann, Jose A. Freixas-Coutin, Jin Seo, Ruth Simmons, Christian Diener, Sean M. Gibbons

## Abstract

The human upper respiratory tract (URT) microbiome, like the gut microbiome, varies across individuals and between health and disease states. However, study-to-study heterogeneity in reported case-control results has made the identification of consistent and generalizable URT-disease associations difficult. In order to address this issue, we assembled 26 independent 16S amplicon sequencing data sets from case-control URT studies, with approximately 2-3 studies per respiratory condition and ten distinct conditions covering common chronic and acute respiratory diseases. We leveraged the healthy control data across studies to investigate URT associations with age, sex and geographic location, in order to isolate these associations from health and disease states. We found several robust genus-level associations, across multiple independent studies, with either health or disease status. We identified disease associations specific to a particular respiratory condition and associations general to all conditions. Ultimately, we reveal robust associations between the URT microbiome, health, and disease, which hold across multiple studies and can help guide follow-up work on potential URT microbiome diagnostics and therapeutics.

## Introduction

The human respiratory system is a complex structure, divided into the upper respiratory tract (URT) and the lower respiratory tract (LRT), and is primarily responsible for exchange of oxygen and carbon dioxide with the atmosphere (1). The respiratory tract, with an approximate surface area of 70 m^2^, is known to harbor a diverse microbial community (2). Beginning at birth, colonization by microbes occurs through constant exposure to the surrounding environment via aspiration (2, 3). A quasi-stable community develops over time, typically consisting of keystone genera such as *Corynebacterium* and *Dolosigrangulum* in young healthy children (4) and *Corynebacterium* and *Staphylococcus* in healthy adults (5). The URT, consisting of the nares, nasal passages, mouth, sinuses, pharynx and larynx, is the section of the respiratory tract most exposed to the environment, and harbors the highest bacterial density (2). Upsetting the balance of this URT microbiome may lead to opportunistic pathogen invasion and serious respiratory tract-related disease and infection (6,7). Chronic respiratory diseases represent the largest disease burden worldwide, affecting over half a billion people in 2017 (8). Pneumonia, an infection of the lungs, is a leading cause of mortality across the world, responsible for an estimated 3.2 million deaths in 2015 (9). The likelihood of being infected by the influenza virus, another common respiratory pathogen that has caused recurrent epidemics over the past century, has been shown to perhaps be dependent on the URT microbiome (7, 10). Additional respiratory conditions, such as RSV, rhinosinusitis, and recurrent respiratory allergies, have all been linked with disruption of the URT microbiome (11–13).

Maintaining a diverse commensal microbiome can be preventative against the invasion of opportunistic pathogens (2, 14). Commensal bacteria can help to saturate metabolic niche space, preventing invasion and engraftment of potential pathogens (6). Additionally, commensals have been shown to directly suppress viral infections through activation of host immune responses (15). Early exposure to commensal microbes can even lead to long-term immunomodulation, preventing autoimmune diseases and promoting tolerance to allergens (16, 17). Overall, the symbiotic relationship between the URT microbiome and the host appears critical for the maintenance of human health (2, 18).

As with the gut microbiome, heterogeneity exists in microbial composition of these communities across individuals. However, several taxonomic patterns of health have been identified in the URT, which are also influenced by variables such as climate, demographics, geographic location, age, and gender. Keystone or core taxa, known to have a general positive association with health, include the genera *Dolosigranulum* and *Corynebacterium* (19–21). For example, the sinonasal area is predominantly colonized by *Corynebacterium* and *Staphylococcus* (*22, 23*), whereas throat and tonsil areas are mostly represented by *Streptococcus*, *Fusobacterium*, and *Prevotella* (*24, 25*). Certain species in the genera *Streptococcus, Haemophilus,* and *Pseudomonas* have been linked to negative health outcomes and disease (1, 19, 26–28). However, respiratory illnesses are often polymicrobial, and are often caused or facilitated by the presence of multiple organisms (29). Identifying consistent signatures of URT health and disease has been hampered by the variability in reported results from individual case-control studies.

Here, we conducted a meta-analysis of the composition of the URT microbiome across health and disease states to identify robust patterns that persist across independent studies within and across multiple respiratory conditions. Using 16S amplicon sequencing data collected from the nasopharynx or the oropharynx across cases and controls from 26 independent studies representing 10 respiratory diseases and conditions, we observe robust associations between the relative abundance of specific genera and disease status. The diseases or conditions included in the meta-analysis are: asthma(30–32), chronic obstructive pulmonary disease (COPD)(33), coronavirus disease 2019 (COVID-19)(34–36), influenza(37–39), pneumonia(14, 40, 41), respiratory allergies(42, 43), rhinosinusitis(44–46), respiratory syncytial virus (RSV)(47–49), respiratory tract infection (RTI, defined as a viral or bacterial infection of the upper or lower respiratory tract, including bronchitis)(50–52), and tonsillitis(53, 54). Knowledge of these associations may help guide the development of diagnostic tools and therapeutic interventions aimed at prevention or treatment of respiratory conditions, by understanding the various types of URT dysbioses that may contribute to these disease states.

## Results

### Assembling case-control studies for a URT meta-analysis

To investigate the associations between the composition of the URT microbiome and disease susceptibility, we analyzed data collected from 26 independent case-control studies including 4701 total samples (**Fig. 1**). Studies included in this meta-analysis had publicly available 16S amplicon sequencing data and associated metadata on disease status. Unfortunately, additional metadata, such as age, gender, and demographic data were not uniformly available across all studies. For each study, the raw data in FASTQ format were downloaded and processed through the same bioinformatic pipeline, defined in the Methods section below. All analyses were conducted at the genus level, given the phylogenetic resolution of partial 16S amplicon sequencing (55). Details on each study included in the meta-analysis can be found in **Table S1**.

**Figure 1.**
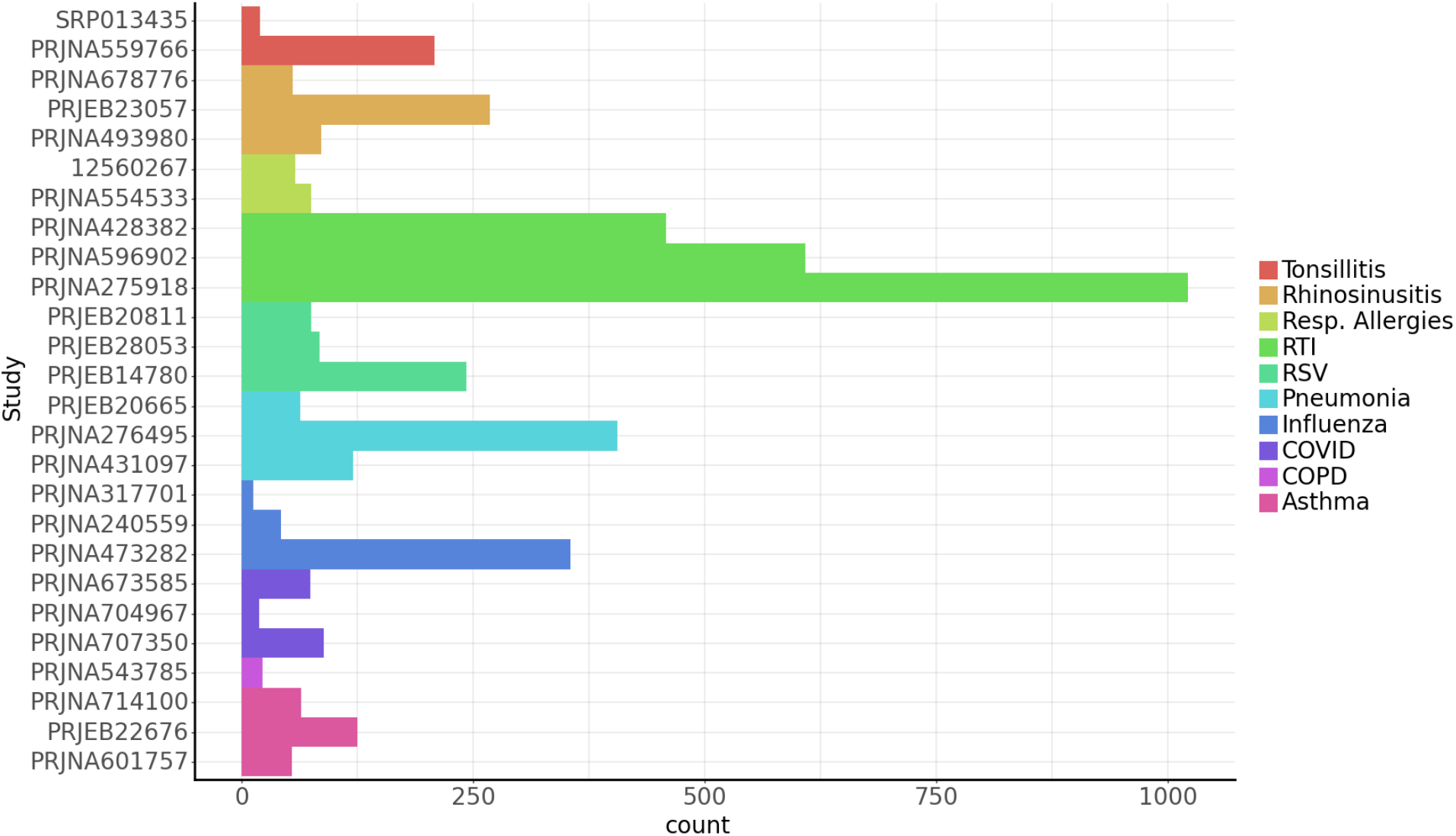
26 independent case-control studies representing 10 respiratory conditions. Studies included between 12 and 1021 subjects (i.e., ‘count’ on the x-axis refers to the number of subjects within a given study). Samples were collected from the nasopharynx or oropharynx, and 16S amplicon sequencing for each URT space was used to analyze ASVs. We attempted to identify three independent studies for each disease condition, but could only obtain 1-2 studies for a handful of conditions.

### Alpha- and beta-diversity analyses show community-wide impacts of disease conditions

We compared URT microbiome alpha-diversity (Shannon index) between disease cases and healthy controls across studies. Shannon diversity was significantly higher in controls compared to cases for several disease conditions, including pneumonia, RSV, respiratory allergies, rhinosinusitis, and tonsillitis, with the exception of influenza, where Shannon diversity was significantly higher in cases compared to controls (**Fig. 2**). It is not well established whether or not alpha-diversity of the URT microbiome is associated with disease (56). These results indicate that a decline in alpha-diversity is frequently an indication of a disease-associated disruption.

**Figure 2.**
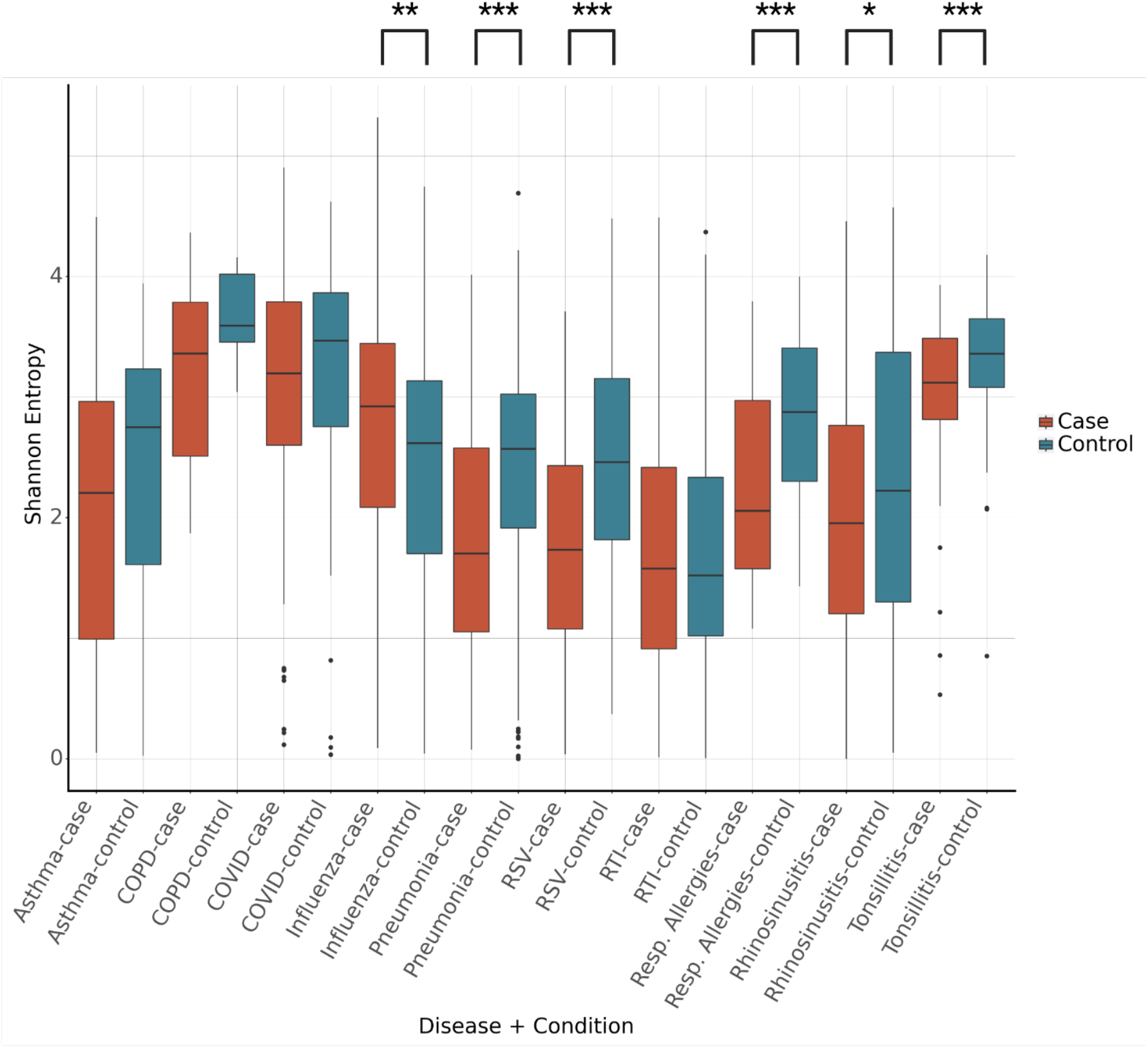
Alpha-diversity between disease cases and healthy controls for each disease condition. The alpha diversity was significantly higher in controls vs. cases for pneumonia, RSV, respiratory allergies, rhinosinusitis, and tonsillitis. The opposite was observed for influenza. Significance between cases and controls determined by independent Student’s t-test, * = p<0.05, ** = p<0.01, *** = p<0.001.

We calculated Bray-Curtis distances at the genus-level, to investigate beta-diversity patterns across studies (**Fig. 3**). Analysis by PERMANOVA showed significant differences in beta-diversity between samples collected from two different URT sites, the nasopharynx and the oropharynx. This is consistent with findings that the nasopharyngeal and oropharyngeal microbiomes are compositionally distinct (57). Additionally, a significant difference was observed between samples taken from different continents, which pushes against prior assertions that the URT microbiome is generally consistent across geographic regions (58). As expected, significant differences were observed in Bray-Curtis dissimilarity in cases relative to controls, as well as between disease conditions (**Fig. 3**).

**Figure 3.**
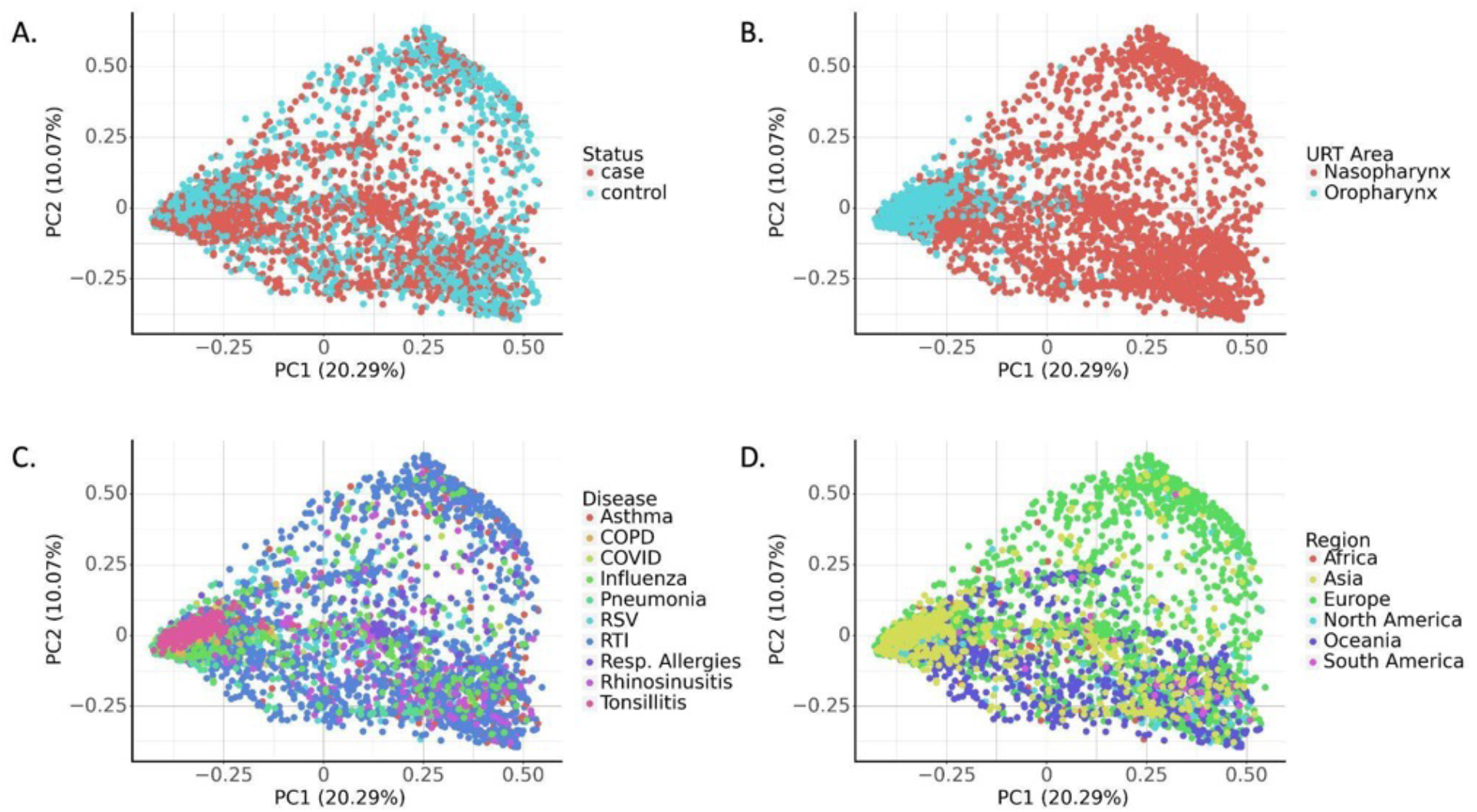
PCoA plots of genus-level Bray-Curtis distances along the first two principal components across the 26 studies URT case-control studies. Beta-diversity was significantly associated with disease status (**A**), URT sampling site (**B**), disease type (**C**), and geographic region (**D**). Significance differences in beta-diversity were observed for all four parameters, as determined by PERMANOVA, *** = p<0.001 in all cases.

### Covariates are significantly associated with URT microbiome composition

Next, we aimed to examine the influence of geographic region, age, and sex on healthy URT microbiome from across studies. Only 38.5% of the studies included metadata on age and sex, which limited our sample size for these variables. Metadata on geographic regions was available for all studies. Kruskal-Wallis one-way analysis of variance was used to determine if a significant difference in centered log-ratio transformed (CLR) relative abundance existed between geographic regions (**Fig. 4**). Stratifying between nasopharyngeal and oropharyngeal samples, results were filtered to those with mean relative abundance greater than 1%. 17 taxa showed a significant enrichment in at least one geographic region (**Fig. 4**). A post-hoc Dunn’s test was conducted on each taxon to determine individual differences (**Table S3**).

**Figure 4.**
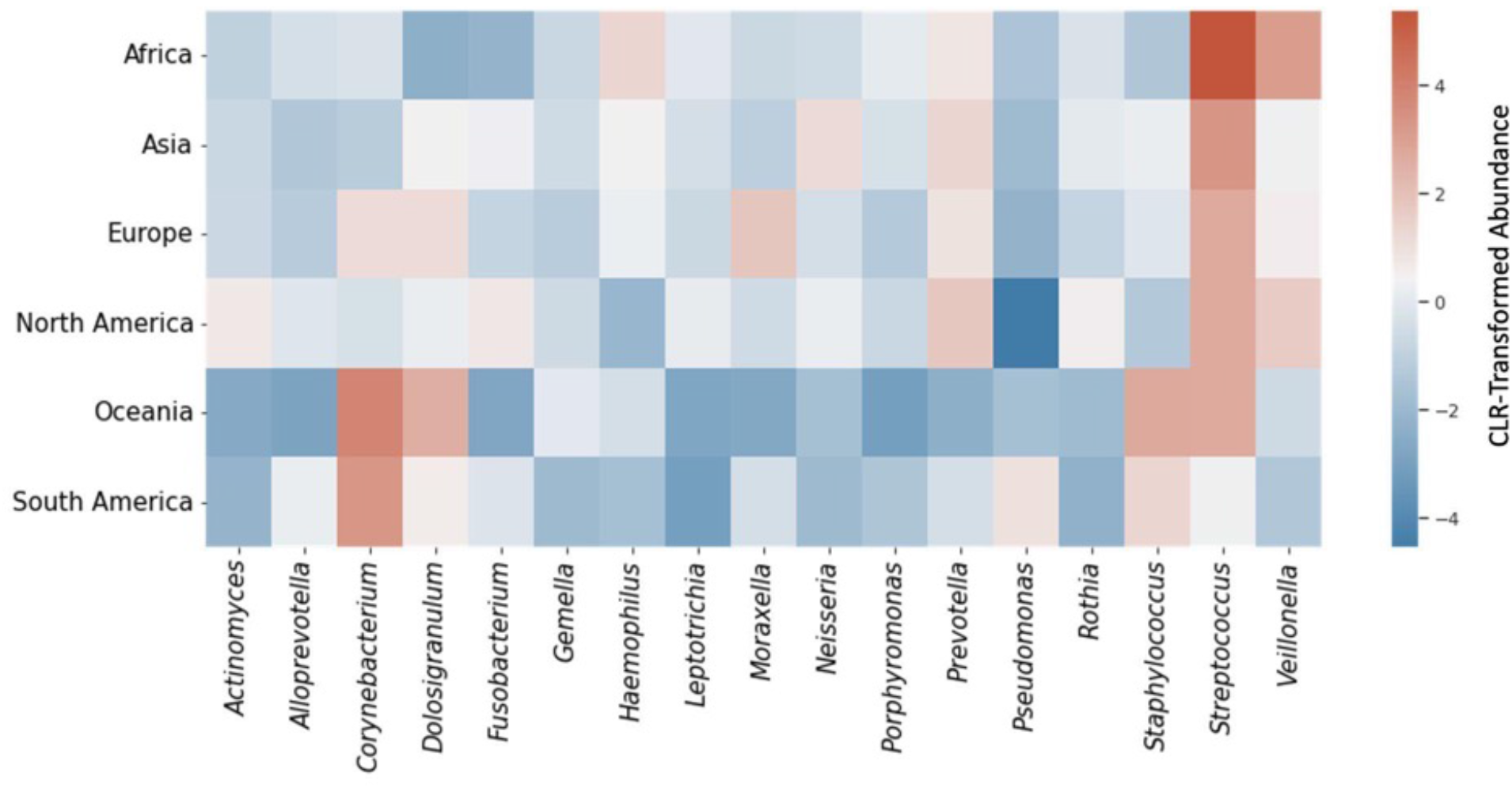
Significant differences in abundance of prevalent taxa between geographic regions in healthy controls. Taxa with significant associations with geographic location are shown. Mean centered log-ratio transformed abundance by geographic region is shown by color encoding, with red indicating higher abundance and blue indicating lower abundance. Significance was determined by Kruskal-Wallis non-parametric analysis of variance, with FDR-corrected p-value < 0.05.

To investigate how relative abundances of URT genera vary with age in healthy populations, regression analysis was conducted, controlling for URT sampling site and geographic location, and treating age as a continuous variable. Overall, 22 taxa were significantly associated with age, based on linear regression containing a squared term for age to uncover non-linear relationships. Samples were grouped into age quantiles, in order to visualize mean CLR transformed abundance across age groups for genera that showed significant associations (**Fig. 5**).

**Figure 5.**
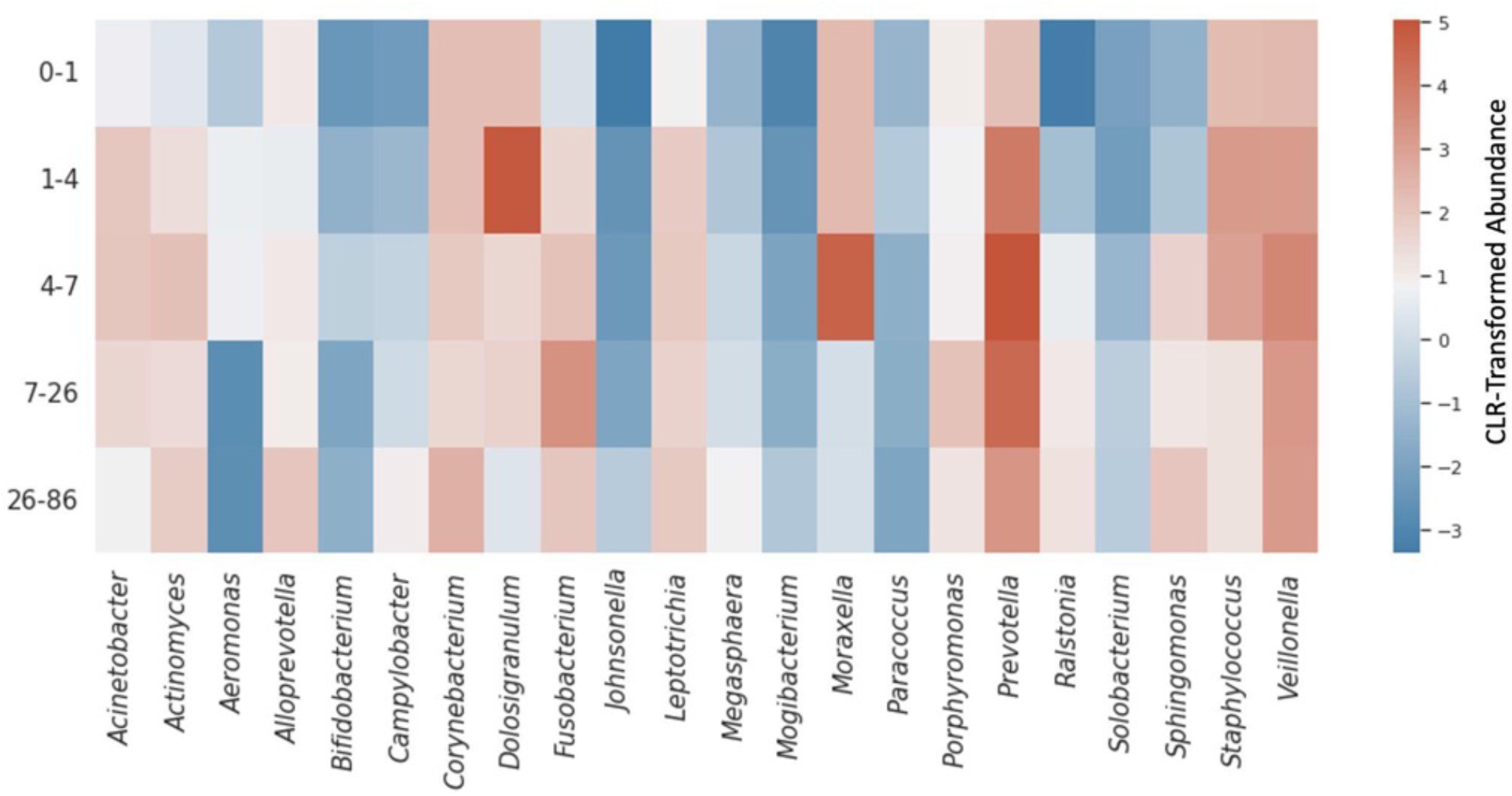
Significant associations in abundance of prevalent taxa between age groups in healthy controls. Taxa with significant associations with age are shown. Mean centered log-ratio transformed abundance by age group is shown by color encoding, with red indicating higher abundance and blue indicating lower abundance. Significance was determined by results of linear least-squares regression, FDR-corrected p-value < 0.05.

Using the same regression framework as in the age analysis, with sex as a categorical variable, no significant associations were observed between sex and microbial abundance in healthy controls, after correcting for URT sampling site and geographic location.

### URT microbiomes show distinct taxonomic associations within and across disease states

We next investigated whether we could identify robust taxonomic patterns of URT microbiome disruption across disease conditions. We conducted logistic regression with disease status as the dependent variable for each genus, including URT sampling site (nasopharynx or oropharynx) and geographic region as covariates. First, we conducted this analysis irrespective of the specific disease condition, across all samples in the meta-analysis, to identify taxa with an overall association with health or disease (**Fig. 6**). Next, we conducted the same analysis between cases and controls for each taxon within each disease condition (**Fig. 6**). Two respiratory conditions, namely COVID-19 and asthma, showed no significant taxonomic enrichments in health or disease (all FDR-corrected p>0.05). Overall, *Acineobacter*, *Haemophilus*, *Pseudomonas*, and *Streptococcus* were generally associated with disease, independent of the specific disease type, while *Alloprevotella*, *Corynebacterium*, *Dolosigranulum, Fusobacterium*, and *Prevotella* were generally associated with health. Many of these same associations were also observed within a given disease type. *Corynebacterium* was associated with health across five disease types (influenza, RTI, respiratory allergies, rhinosinusitis, and tonsillitis). Conversely, *Acinetobacter* and *Streptococcus* were associated with disease across three disease types (influenza, pneumonia and respiratory allergies for *Acinetobacter*; influenza, pneumonia and RTI for *Streptococcus*; **Fig. 6**). However, not all associations were consistent across disease contexts. For example, *Cornyebacterium* was significantly associated with disease in RSV, and *Pseudomonas* was significantly associated with health in RSV. Pneumonia and RTI showed the largest number of significant associations, each with 11 associations with either health or disease status (**Fig. 6**). *Streptococcus* had the highest mean relative abundance of taxa with significant associations, at 17.2% ± 0.3%, followed by *Corynebacterium*, *Staphylococcus*, *Dolosigranulum*, *Haemophilis*, and *Prevotella*, all with mean relative abundances over 5% (**Fig. 6**). Effect size and *p* value were recorded for each taxon-disease pair (**Table S4**).

**Figure 6.**
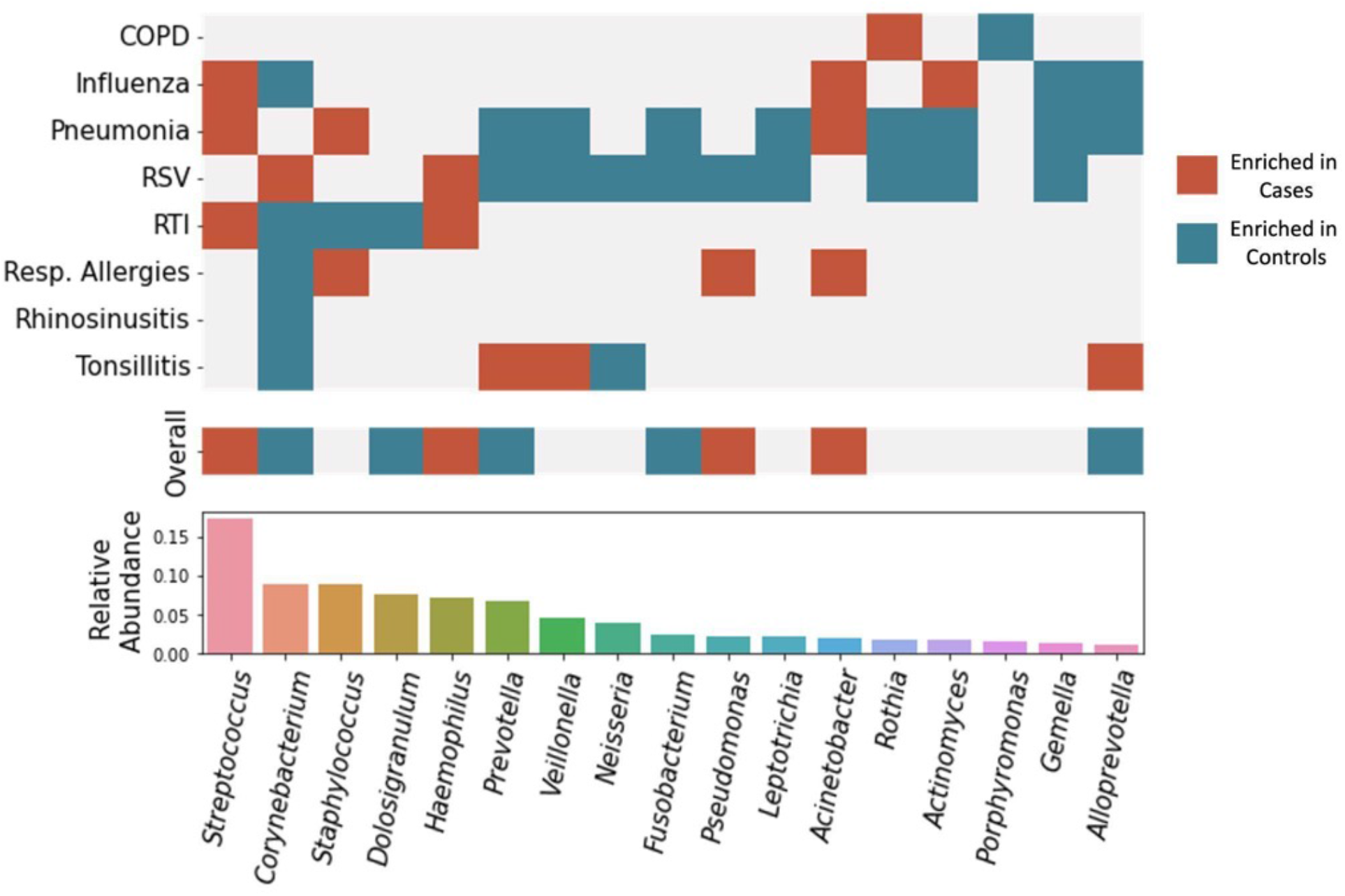
Logistic regression results at the genus-level, controlling for sampling site and geography, within and across diseases. Taxa enriched in cases are denoted in red, and those enriched in controls are denoted in blue. Blank spaces indicate no significant association. Only taxa with at least one significant association are shown. The ‘Overall’ bar shows associations that hold across all samples from all disease types. Mean relative abundance of included taxa among all samples in the meta-analysis are shown in the barplot in the bottom panel. Significant associations defined by the results of logistic regression, FDR-corrected p-value < 0.05.

## Discussion

The results of this meta-analysis were consistent with prior findings regarding the composition of the URT microbiome in health and disease (1), and revealed novel compositional patterns within and across diseases and between healthy individuals across age and geography. They also underscore the importance of recognizing different types of dysbioses in the URT microbiome that can potentially contribute to disease.

Observations of URT microbiome samples showed a significant trend toward lower alpha-diversity in disease cases as opposed to healthy controls for tonsillitis, rhinosinusitis, respiratory allergies, RSV and pneumonia (**Fig. 2**). Previous studies have reported similar signatures in cases of bacterial or viral infection (59, 60). However, conflicting findings suggest that alpha-diversity patterns vary depending on the disease context (61). We found that a decrease in the abundance of keystone taxa associated with health, such as *Dolosigranulum* or *Corynebacterium*, was associated with a decline in alpha-diversity for several disease conditions. Respiratory allergies also showed a significant decrease in alpha-diversity, which could be due to similar perturbations of similar keystone taxa due to host inflammation (2, 12). Influenza was the sole respiratory condition that showed significantly higher alpha-diversity in disease cases. However, this finding will need further validation, as prior reports have found no association between URT alpha-diversity and susceptibility to influenza infection (7, 62).

Bray-Curtis dissimilarity between URT communities were associated with multiple covariates: case versus control status, sampling site (nasopharynx or oropharynx), disease type, and geographic region (**Fig. 3**). Concordantly, prior work has shown significant beta-diversity differences between health and disease states (59) and separation between nasopharyngeal and oropharyngeal samples, with the oropharynx harboring a more diverse microbial population than the nasopharynx (39) (**Fig. 3**). Finally, significant beta-diversity differences between samples from distinct geographic regions were novel but somewhat expected, despite prior assertions of a lack of geographic signal (58), as microbial colonization is dependent on interactions with the surrounding environment (**Fig. 3**). Sex and age metadata were only available in a minority of the studies included in the meta-analysis, limiting our capacity to investigate how these variables shaped alpha- and beta-diversity. However, we did explore specific taxonomic associations with these variables within healthy control samples, as we discuss below.

We next looked into how covariates including age, sex and geographic location shaped the taxonomic composition of the URT microbiome in healthy individuals across studies, in order to separate these signals from health and disease associations, and indicate which covariates should be considered in future analyses. For example, *Streptococcus*, which has been associated with disease in prior studies, was observed at significantly higher abundance in Africa and Asia (**Fig. 4**). Thus, the development of robust diagnostic and therapeutic tools must take these geographic signatures into account. We included geographic region as a covariate in our disease regressions below in order to account for this potentially confounding heterogeneity. Age also proves to display significant associations with composition. Two keystone taxa, *Dolosigranulum* and *Moraxella*, were enriched in children as compared to adults, as previously reported (4), (^63^) (**Fig. 5**). Additionally, we saw an increase in the keystone taxon *Corynebacterium* in adults, when compared to children (**Fig. 5**). Due to the breadth of associations observed with age, and the purported inhibition of pathogenic invasion by some of these age-associated genera (2), we suggest that age should be included as a covariate when analyzing URT microbiome data, whenever possible. However, when age metadata are unavailable, we hope that the list of taxa provided here can be used to identify associations that may be confounded by age. We saw no significant associations with sex in healthy controls, even after correcting for URT sampling site and geography.

Robust taxonomic signatures associated with case-control status across studies were observed independent of URT sampling site and geographic region (**Fig. 6**). Across all samples, irrespective of disease type, significant signatures were observed for several keystone taxa that tend to be enriched in healthy individuals (2), like *Alloprevotella, Corynebacterium, Dolosigranulum, Fusobacterium* and *Prevotella*. Of these, *Corynebacterium, Dolosigranulum, Fusobacterium* and *Prevotella* have been previously identified as core taxa, putatively associated with health (1). Additionally, these are largely abundant taxa, with mean relative abundance above 5% for *Corynebacterium, Dolosigranulum* and *Prevotella* (**Fig. 6**). Conversely, several genera known to harbor opportunistic pathogens, including *Acinetobacter, Haemophilus, Pseudomonas* and *Streptococcus*, showed associations with disease, independent of disease type. *Acinetobacter baumannii, Haemophilus influenzae, Pseudomonas aeruginosa* and Group A *Streptococcus* are all well established to cause disease in humans (26–28, 64). We also observed several disease-type-specific signatures. For example, pneumonia and RSV both showed 11 associations, of which 8 and 9 were depleted in the disease condition, respectively (**Fig. 6**). This supports our earlier hypothesis that lower alpha-diversity in pneumonia and RSV cases versus controls resulted from depletion of beneficial, health-associated taxa. Three taxa, *Acinetobacter, Actinomyces,* and *Streptococcus*, were enriched in individuals with influenza (**Fig. 6**). Invasion of these taxa into the URT microbiome may increase susceptibility to influenza, hence explaining the increased alpha-diversity observed in influenza cases (**Fig. 3**). Interestingly, *Streptococcus* was the taxon with the highest mean relative abundance found to be disease-associated, with specific associations with RTI, pneumonia and influenza. *Streptococcus, Haemophilus* and *Acinetobacter* showed exclusive disease-enrichments (i.e., never enriched in controls), indicating that these taxa may be disease-indicators and potential targets for antimicrobials (26, 28, 64). Conversely, those taxa associated only with health may be good targets for probiotic development, which has shown recent promise in the URT (1). While *Corynebacterium* was enriched in healthy controls in influenza, RTI, respiratory allergies, rhinosinusitis, and tonsillitis studies, it was associated with disease across RSV studies (**Fig. 6**). Here we see another example of dysbiosis taking many forms, and the health or disease associations of many taxa showing strong context-specificity. Understanding which taxa are strongly related to health in which contexts, will further aid the development of diagnostics and therapeutics.

Overall, these findings point to different flavors of dysbiosis that distinguish different disease states in the URT. In some cases, the disease state is characterized by a loss of putatively beneficial commensals, and in other cases it is characterized by the gain of putatively pathogenic taxa, which mirrors what has been seen across diseases in the human gut microbiome (65). The associations identified in this meta-analysis were independent of several important covariates and held true across multiple independent studies. Future work should leverage these results to help guide the development of diagnostics and therapeutics for the URT.

## Materials and Methods

### 16S amplicon sequencing URT cohorts

All phylogenetic and abundance data used in this study consisted of 16S amplicon sequencing data, with multiple hypervariable regions sequenced across studies, spanning the V1 to V7 regions. In total, data from 26 case-control clinical studies were included in the meta-analysis, covering 10 URT-related conditions (asthma, chronic obstructive pulmonary disease, COVID-19, influenza, pneumonia, respiratory allergies, rhinosinusitis, RSV, respiratory tract infection, tonsillitis). A full list of studies, along with links to SRA accession numbers and accompanying metadata, can be found in Supplemental Table 1. The studies contained between 12 and 1,021 subjects, and varied in age from birth to 85 years old. Studies were conducted in all six inhabited continents, with more representation from Europe and North America. With exception of one study for which data was not yet publicly available, 16S sequencing data consisting of FASTQ files for all samples per study as well as any associated metadata for all studies, were downloaded from the NCBI SRA. While some studies included paired-end sequencing reads, only forward reads were used to maintain better analytical consistency across all studies. Publicly unavailable data for one study were obtained via FigShare after direct correspondence and agreement with the author (42). Following data collection, all FASTQ data were imported into QIIME2 for further processing and analysis. Data were imported through construction of a single-end Phred33v2 FASTQ manifest for each dataset. Following import, quality control and filtering in the QIIME2 DADA2 plug-in removed chimeric sequences, trimmed left ends of all sequences by 10bp to remove primers, truncated sequences uniformly at 200bp, and identified amplicon sequence variants (ASVs).

### Data preprocessing and taxonomic classification

The Silva high quality rRNA gene database version 138 was used to assign taxonomy to ASVs (66). The full-length 16S classifier was used due to heterogeneity in the hypervariable region used for sequencing between studies. Mean classification at the genus level was 74.3% of reads (Table S2; Fig. S1). At the species level, classification was less successful, with mean classification of 9.3%. As a result, all subsequent analyses were conducted at the genus level by binning ASV counts together based on their genus-level annotations.

### Alpha diversity analyses

To investigate alpha-diversity, QIIME2 artifacts containing sequences for each study were merged into a single dataframe. Prior to calculation, algorithmic filtering removed any taxa with fewer than two reads per study, and any taxa present in less than 50% of samples across a study. Rarefaction was conducted to a sampling depth of 2000. This merged data frame was converted into a QIIME2 artifact, and diversity metrics were calculated in QIIME2. Shannon entropy was used to estimate alpha-diversity for all samples included in the meta-analysis. Differences in Shannon entropy were explored across metadata parameters. Of particular interest was the difference in alpha-diversity between cases and controls. Shannon entropy for cases and controls within each disease were plotted and tested for significant differences using a two-way ANOVA (p<0.05) in SciPy (v1.8.1).

### Beta-diversity analyses

To estimate beta-diversity, the filtered and rarefied table constructed previously was used to construct a Bray-Curtis dissimilarity matrix using Scikit-Bio (v0.5.7). Subsequently, PCoA was used to analyze and visualize overall beta diversity within Scikit-Learn. Significant differences in beta diversity were observed along multiple axes, including case vs. control status, disease type, geographic location and URT sampling site, as determined by PERMANOVA in Sckit-Bio (p<0.05).

### URT compositional patterns across geographic regions

A genus-level abundance matrix was constructed using only healthy control samples, and taxa with fewer than two reads per study or those present in fewer than 50% of samples across a study were removed. To examine the association between geographic location and centered log-ratio (CLR) transformed relative abundance of common taxa, Kruskal-Wallis one-way analysis of variance was used to determine significant enrichments of taxa in each geographic region using SciPy. Samples were stratified into oropharyngeal and nasopharyngeal samples, and results from each were combined into the final result. For the purpose of these analyses, continents in which studies took place were used as the geographic regions. As sex and age metadata were not available for 60% of the studies, these covariates were not accounted for in this analysis. The p-values, as determined by Kruskal-Wallis, were corrected for multiple comparisons using the Benjamini-Hochberg FDR correction procedure (67). Taxa identified to be significantly enriched in one geographic region (FDR-corrected p<0.05) were added to a heatmap, with color encoding the average CLR transformed relative abundances in each context. Post-hoc Dunn’s test was conducted to determine significant enrichments for each taxon in each geographic region, using Scikit-posthocs (v0.7.0) (68).

### URT microbiome-age associations

Associations between age and CLR transformed relative abundances was analyzed via multiple regression in *statsmodels* (v0.13.5)(69), as were all regressions in this analysis. Using 10 studies for which age metadata was available, regressions were conducted using the following formula “clr ∼ age + age^2^ + URT_site + region” that was used to determine significant enrichments across age, accounting for URT sampling site and geographic region. The square term for age was included to determine if non-linear relationships existed between CLR and age. The p-values were corrected for multiple comparisons via Benjamini-Hochberg FDR correction as previously described. Significantly associated taxa (FDR-corrected p<0.05) were added to a heatmap with color encoding CLR transformed relative abundances as before.

### URT microbiome associations with sex

Associations between sex and relative abundance were determined via multiple regression. Using the 10 studies for which sex metadata was available, regressions were conducted using the following formula: “clr ∼ sex + URT_site + region”. After FDR correction, no significant associations remained.

### URT microbiome-disease associations

To investigate the association between URT genera and disease, sample read counts were normalized using a CLR transformation. Logistic regressions used case-control status as the dependent variable and CLR-transformed abundance, URT sampling site, and geographic location as independent variables, following the formula “case_control_status ∼ clr_genus + URT_site + region”. Next, the same analysis was conducted for samples within each disease group, to determine disease-specific associations. Geographic regions were not sufficiently represented within individual disease conditions, hence geography was dropped as a covariate in the disease-specific analyses. Thus, the formula “case_control_status ∼ clr_genus + URT_site” was used for disease-specific regressions. Mean relative abundance of each taxon found to be significant was calculated for visualizations. The Benjamini–Hochberg procedure for adjusting the False Discovery Rate (FDR) (67) was performed to account for multiple testing. Significance was assigned to any association with an FDR-corrected p-value less than 0.05. Results were plotted in a binary heatmap, with significantly health-associated values designated in blue and disease-associated values designated in red. Heatmaps were constructed using Seaborn (v0.12.2).

### Data and code availability

Information on accessing the raw data and metadata from all 26 case-control studies included in this analysis can be found in **Table S1**. Analysis code and intermediate data files can be found at the following GitHub repository: https://github.com/Gibbons-Lab/2023_URTmetaanalysis

## Acknowledgements

This work was supported by a research grant from Reckitt Health US LLC to SMG. This work was also supported by a Washington Research Foundation Distinguished Investigator Award and by startup funds from the Institute for Systems Biology to SMG.

## Supplemental Figure Caption

**Figure S1.**
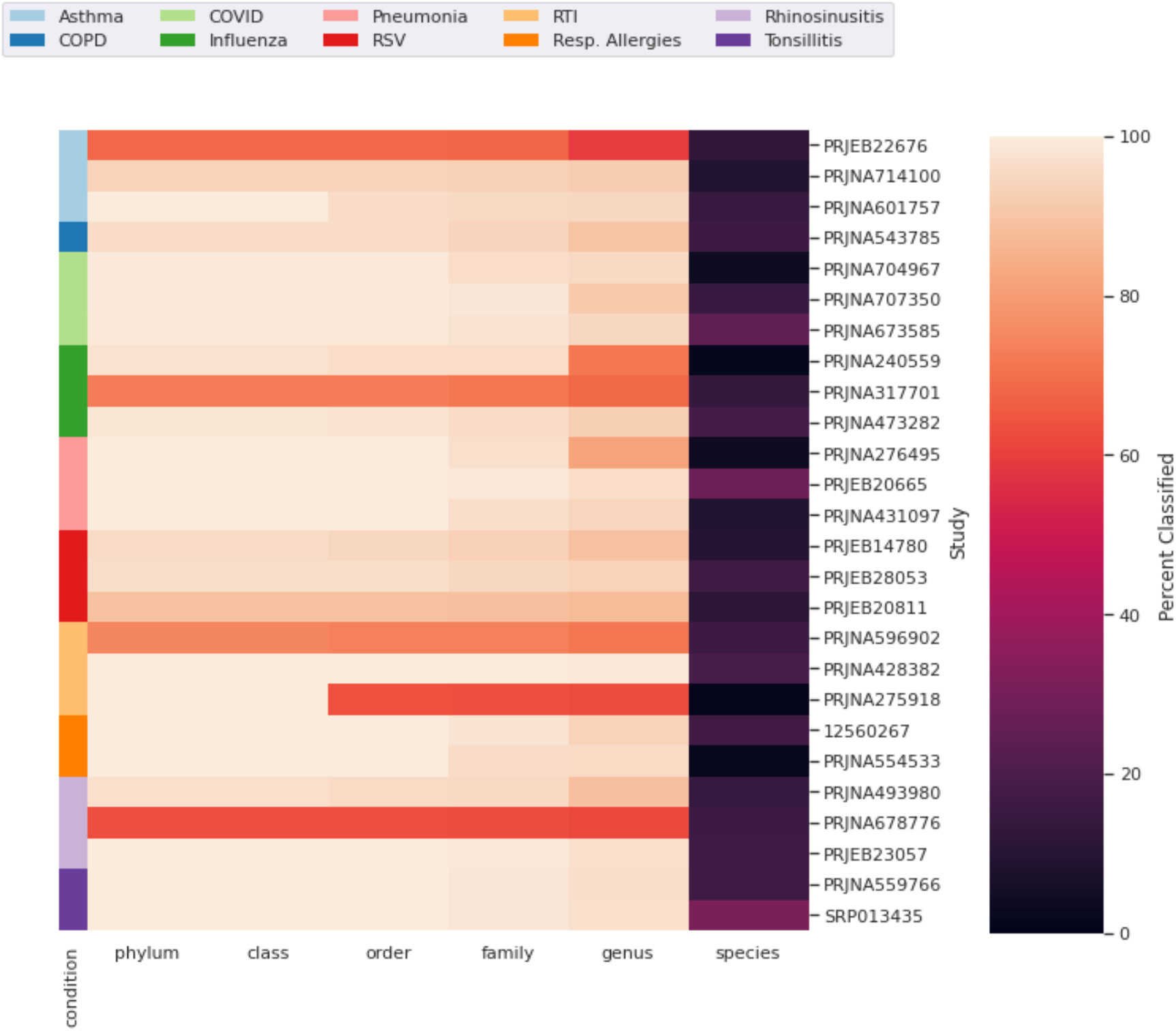
Mean classification percentage for each study at each taxonomic level. Classification remained at or above 60% for all studies through the genus level. At the species level, a significant drop in classification was observed.

